# Genetic targeting of astrocytes associated with specific neuronal circuit in adult *Drosophila*

**DOI:** 10.1101/2025.02.12.637801

**Authors:** Joana Dopp, Frederik Hobin, Sofia Mastroianni, Jiekun Yan, Lisa van Ninhuys, Sha Liu

## Abstract

Astrocytes are the major glial population of the brain and have been associated with a vast number of functions. To probe this diversity and to reach a similar level of understanding about astrocyte physiology that we have about neurons, we need genetic tools to target specific astrocytic subpopulations. In *Drosophila*, we are restricted to using driver lines that drive expression in astrocytes throughout the brain. To target specific astrocytes, we have optimized the genetic tool TRACT (and refer to it as astro-TRACT), allowing effector expression specifically in local astrocytes of a given neuronal circuit. We analyzed specificity, sensitivity and reproducibility of the tool across various MB split-Gal4 drivers. We found that the number of pre-synapses correlates positively with the success of the tool. Applying the tool to characterize morphology of individual astrocytes revealed that local astrocytes around MB medial compartments project into the ellipsoid body. Astro-TRACT will be a valuable resource to investigate both mechanistic astrocyte-neuron signaling and functional and structural astrocytic diversity across the adult *Drosophila* brain.

## Introduction

Genetic tools have advanced our understanding of brain function and connectivity. In *Drosophila*, tools that facilitate labeling and manipulating specific cellular subpopulations are necessary to address mechanistic neuroscientific questions [1]. Together with neurons, glial cells populate the brain, and astrocytes make up their largest class. In the past, astrocytes have been viewed as housekeeping cells. In contrast, astrocytes are active participants in many physiological processes such as synaptic plasticity [2], gliotransmission of ATP, cytokine signalling, recycling and buffering of neurotransmitters and ions [3] and downstream behaviours such as sleep homeostasis [4]. Specific astrocytes involved in regulating a given process may be located close to a relevant neuronal circuit. Understanding how such local astrocytes regulate processes requires functional studies.

In *Drosophila* neurons, genetic control of specific cell populations has been achieved by the generation of an extensive library of thousands of driver lines targeting specific neuronal subpopulations [5,6]. Unfortunately, a similar feat appears unrealistic for glial cells. In contrast to mammalian astrocytes, transcriptionally there are little differences between fly astrocytes located in different regions of the brain [7]. Furthermore, the synthetic Gal4 collection covering transcriptional enhancers across the fly genome [5] does not contain any region-specific astrocyte promoters [8]. Despite the lack of transcriptional diversity, the morphological diversity of astrocytes across the fly brain is well-documented [7,8]. For example, astrocytes around the Mushroom Body (MB) have unusual thin branches [8]. The region-specific glial structural patterns have been attributed to specialized, dynamic subtypes that adapt in response to cues from their local environment [7,8], and may serve different functions in local circuits.

Here, we describe another route to achieve targeting of local astrocyte subpopulations, that makes use of the extensive library of neuronal drivers instead of relying on genetic differences between astrocyte populations. We adapted the TRACT (<u>Tra</u>nsneuronal <u>C</u>ontrol of <u>T</u>ranscription) system, which is based on synthetic Notch-mediated transmembrane proteolysis [9,10]. It was initially developed to study neuronal wiring and cell-cell interactions, including glia-neuron contacts in the developing fruit fly. Huang et al. (2016) characterized glial expression across multiple larval brain regions and compared the expression between *alrm* and *repo* enhancers. Here, we have validated and optimised the TRACT tool for adult flies, analyzed its specificity, sensitivity and reproducibility and applied it to investigate morphological patterns of MB – associated astrocytes. We found that astrocytic labelling is achieved only when sufficient synapses are present in their associated MB compartment. Additionally, we found that local astrocytes of MB γ, β and β’ compartments connect to the EB.

## Results

### Flexibility of astro-TRACT is increased by using split-Gal4 system to control specific neuronal circuits

TRACT involves the expression of at least four transgenes in two binary systems, two of which are driving expression in a chosen neuronal circuit and the other two driving expression in astrocytes. The specificity of TRACT is determined by the specificity of the neuronal driver. To allow for high flexibility of choosing neuronal drivers, we developed TRACT in different combinations of binary systems, most notably to use the Gal4>UAS system for defining the neuronal circuit, while using the QF2>QUAS system to drive effector expression in astrocytes (**Fig. S1**). Reserving the Gal4>UAS system for the neuronal part is advantageous because the highest specificity of neuronal drivers is achieved with split-Gal4 drivers. Split-Gal4 drivers consist of two parts - VP16AD and Gal4DBD, that will form a functional Gal4 protein only when together and consequently only in cells that express both enhancers. In addition to the extensive split-Gal4 library that others have generated [6], any combination of VP16AD and Gal4DBD lines can be recombined to achieve targeting of highly specific populations. Such specific drivers can subsequently be crossed to TRACT to examine astrocytes close to those populations. Labelling intensity and expression pattern is similar for different transcription factors (Gal4 or QF2) being used to drive reporter expression (**Fig. 1c-e, Fig. S2**). We observed labelling differences rather across injection sites and astrocyte enhancers (*alrm* and R86E01).

**Figure 1.**
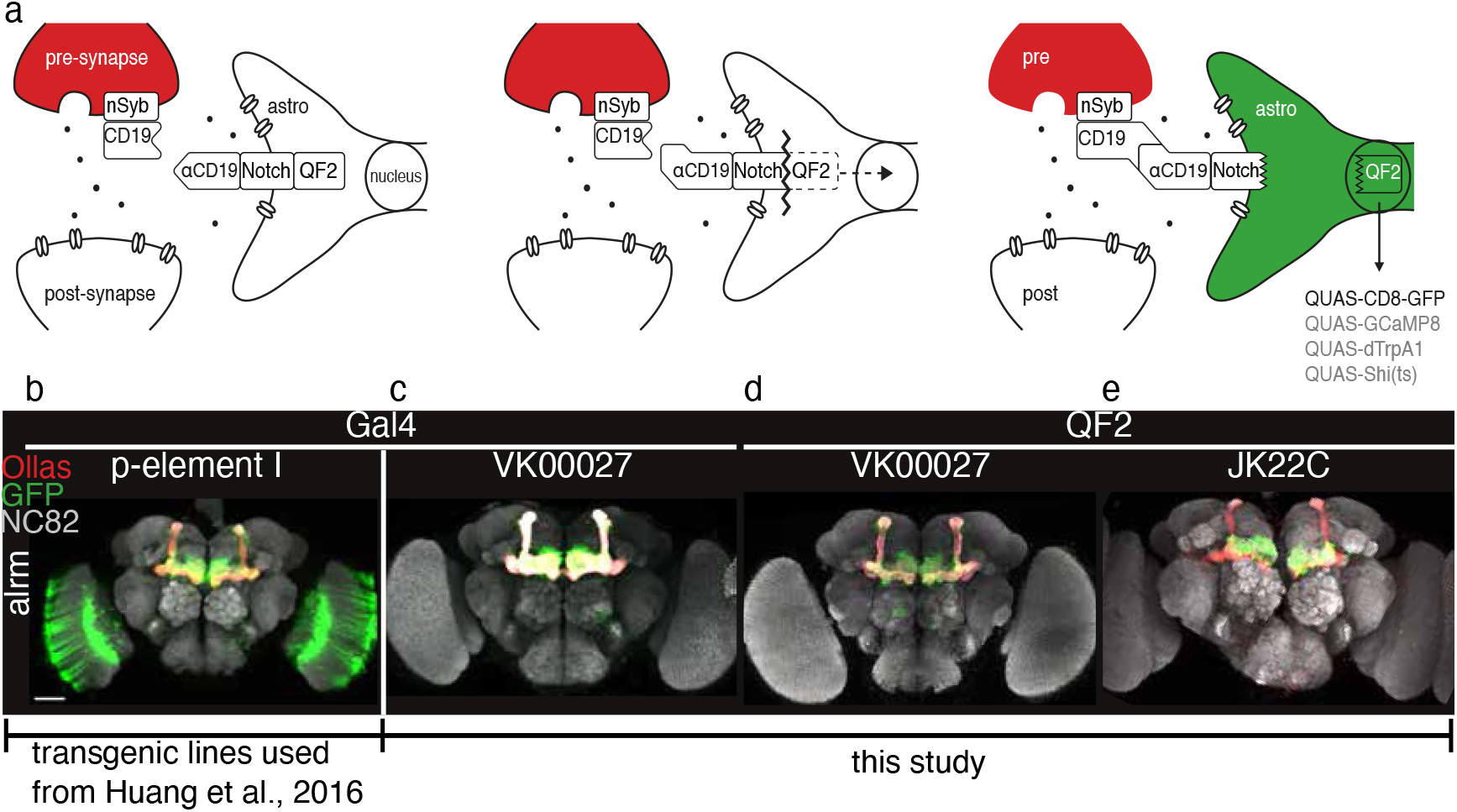
Notch-mediated genetic tool labels local astrocytes with high precision. **a**. Schematic representation of sequence of events occuring with G-CLIP tool. Left: Extracellular CD19 ligand at the neuronal pre-synaptic site connects to extracellular anti-CD19 domain fused with a modified Notch receptor, which is located in the membrane of all astrocytes. Middle: The transcription factor QF2 is cleaved from the modified Notch receptor and translocates into the nucleus. Right: Once in the nucleus, QF2 will bind to QUAS, thereby expressing green fluorescence in local astrocytes. Any effector in place of GFP can be used. The tool contains four genetic elements: (1) split Gal4 driver, (2) UAS-nSyb-CD19-Ollas, (3) R86E01/alrm-DSCP-nlg-synNotch-anti-CD19-QF2, (4) QUAS-effector. **b**. Astrocyte labelling around the Mushroom Body (MB) with original version by Huang et al., 2016. Scale bar 50 μm. **c-e**. Astrocyte labelling around MB with (c) Gal4 or (d-e) QF2, injected in either (c-d) VK00027 or (e) JK22C with alrm enhancer.

While in the original version by others [9] the astrocyte Gal4 transgenic element was inserted into the second chromosome by p-element, we injected either into VK00027 (third chromosome) or JK22C (second chromosome) landing sites and achieved higher specificity of astrocyte signal around the MB (**Fig. 1b-e)**. We did observe unspecific neuronal labelling in the posterior part of the brain with the R86E01 - Gal4 version injected into VK00027, but not with the *alrm* enhancer. Therefore, we did not inject the R86E01 - QF2 version into VK00027, but only into JK22C (**Fig. S2b**). Typically, we found that the enhancer R86E01 returns a stronger GFP signal compared to *alrm* (**Fig. 1c-e, Fig. S2**).

### Sensitivity of astro-TRACT is driven by the number of pre-synaptic sites in targeted brain region

Beside testing the specificity of the tool, we asked how sensitive and reproducible the astrocyte labelling is. We tested astrocyte labelling with 18 split-Gal4 drivers that express neurons with their pre-synaptic sites in different parts of the MB lobe compartments. 14 of the 18 drivers label at least one astrocyte. Importantly, the labelling is reproducible across brains of the same genotype (**Table S1**).

While all 18 drivers project axons into MB lobe compartments, they drive effector expression in different cell types: PAM and PPL1 dopaminergic neurons, KCs, and MBONs. We found that successful astrocyte labelling depends on the chosen associated neuronal cell type. While all drivers targeting PAM dopaminergic neurons and KCs were successful, all drivers targeting PPL1 dopaminergic neurons and MBONs were not. We hypothesized that the reason for this discrepancy may be that the number of pre-synapses in a given MB compartment to catalyze the Notch mechanism of TRACT differs between cell types. To test this quantitatively, we estimated the number of pre-synapses per MB compartment based on the public hemibrain connectome [11,12] detailing synaptic connectivity across the fly brain. Indeed, those neuronal driver lines that result in labelled astrocytes typically have a higher number of pre-synaptic sites in MB lobe compartments (**Fig. 2m, Table S2**). Our analysis indicates that around 750 pre-synapses are required in an MB compartment to catalyze the TRACT mechanism and label local astrocytes (**Fig. 2m, Table S2**). In the future, the accuracy of this analysis can be improved by increasing the number of cells matched to drivers as more traced neurons become available in the connectome. Still, estimating synaptic density may be a useful initial predictor of astro-TRACT’s sensitivity in different neuronal circuits.

**Figure 2.**
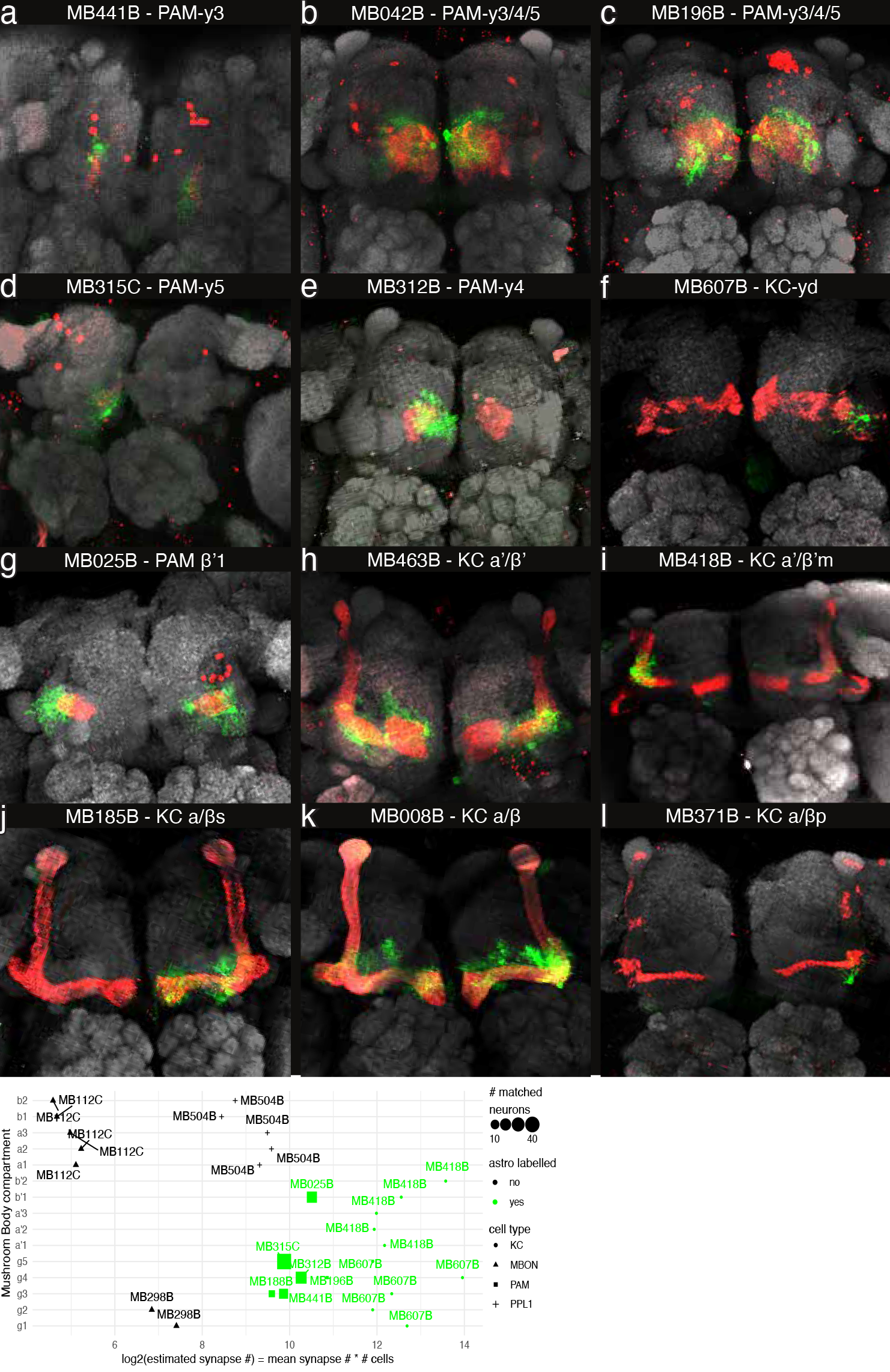
Astrocyte labelling around different compartments of the Mushroom Body. a-f. y lobe g-i. a/b' j-l. a/b. Scale bar 50 um. j. Comparison of pre-synapse number across MB driver lines targeting KC, MBON, PAM, PPL1 neurons, used in this study.

To probe the sensitivity of astro-TRACT in other circuits, we tested two different drivers targeting EB-R5 neurons. Similar to other sparse cell populations (MBON and PPL1 in **Table S1**), we found that the labelling was unsuccessful for one driver of R5 cells (**Fig. S2b**).

Interestingly, we did observe consistent labelling when using the other R5 driver (**Fig. S2a**), suggesting that slight differences in the targeted subpopulation of R5 cells can determine the result.

### Local astrocytes around Mushroom Body connect to Ellipsoid Body

Next, we were interested in applying the tool to examine individual astrocyte morphology around the MB and asked whether MB-local astrocytes project into the Ellipsoid Body (EB). The EB is situated directly posterior to the MB, yet according to the Drosophila connectome there are no neuronal connections between these structures [13-15]. Even more than their physical proximity, this is surprising because they share similar functions. For example, both EB and MB are involved in regulating sleep behaviour.

We asked whether instead of neuronal connections, astrocytes indirectly connect the MB to the EB. To address this question, we combined the optimized TRACT tool with a multi-colour mosaic system. MCFO (MultiColor FlpOut) is a technique to stochastically label individual cells with three tags [16]. The combination of TRACT and MCFO allowed us to visualize individual astrocytes that are located specifically around the MB (**Fig. 3**). The MB-local astrocytes do project into the EB ring. The combination of the three tags of the MCFO tool covers almost the entire circular structure of the EB (**Fig. 3**).

**Figure 3.**
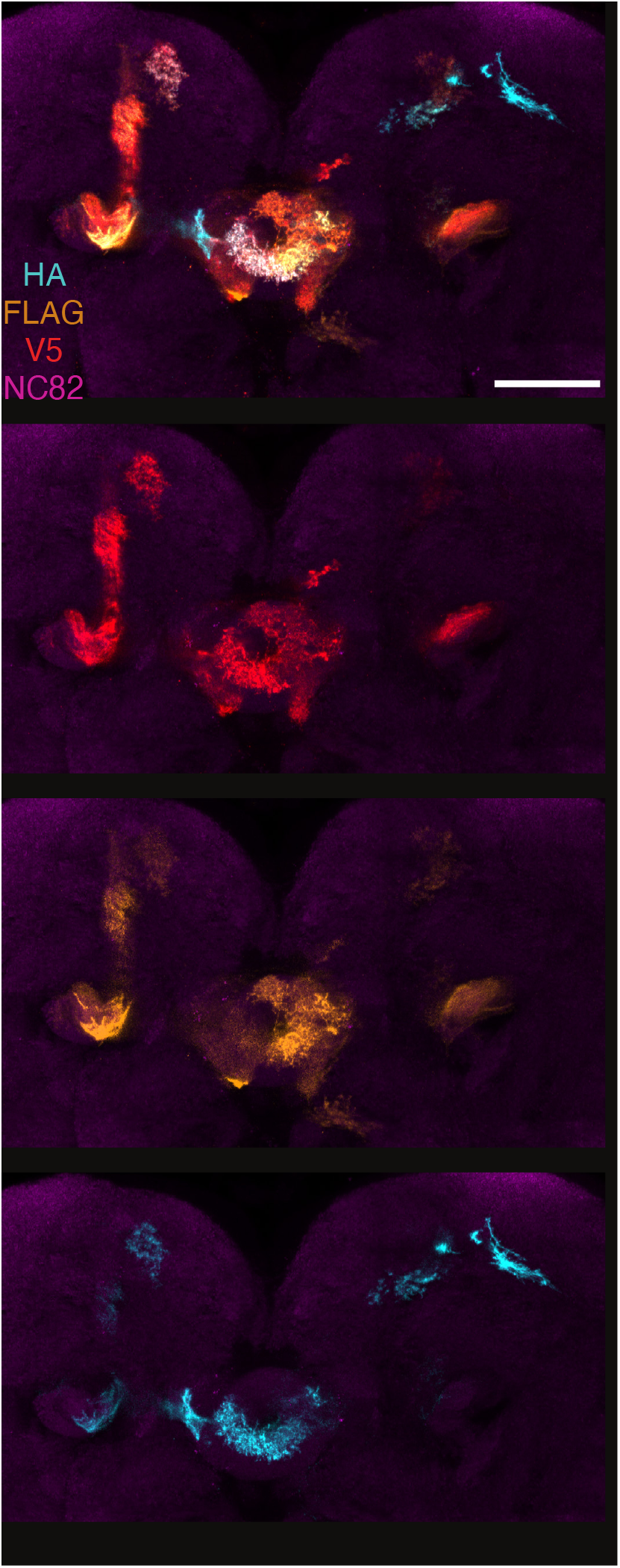
Local astrocytes of Mushroom Body connect to Ellipsoid Body. Composite image (top) and separated channels. Single channel images show labelling of individual astrocytes. Maximum intensity projections include all z planes with the EB structure present. Scale bar 50 μm.

### Astrocytes specifically bridge medial MB compartments of the horizontal lobes with the EB

Next, we asked whether the MB-astrocyte-EB connection is specific to certain MB compartments. Probing this connection required an increased resolution to visualize individual astrocytes. For this purpose, we analysed single astrocyte labelling with TRACT across 14 split-Gal4 drivers targeting the pre-synapses in different MB compartments (**Table S1**).

We found that astrocytes connect a respective MB compartment to the EB for 6 of the 14 driver lines tested (**Table S1, Figs. 3, 4b-d, Video S1**). Typically, astrocytes connecting MB and EB are associated with β’2, β2, γ4 and γ5 and notably often cover more than one compartment at the same time. Interestingly, these astrocytes are not limited to the α/β lobe located closest to the EB, but they extend to the anterior γ lobe (**Table S1**). The astrocytes labelled with the remaining drivers that do not connect to the EB are not located medially but on the lateral side of the MB lobes. We also observed that astrocytes typically project processes within the same hemisphere of their soma’s location, except for two PAM neuron-specific drivers labelling the γ3, γ4 and γ5 MB compartments (MB196B and MB042B), in which cases the astrocytes span across the midline.

**Figure 4.**
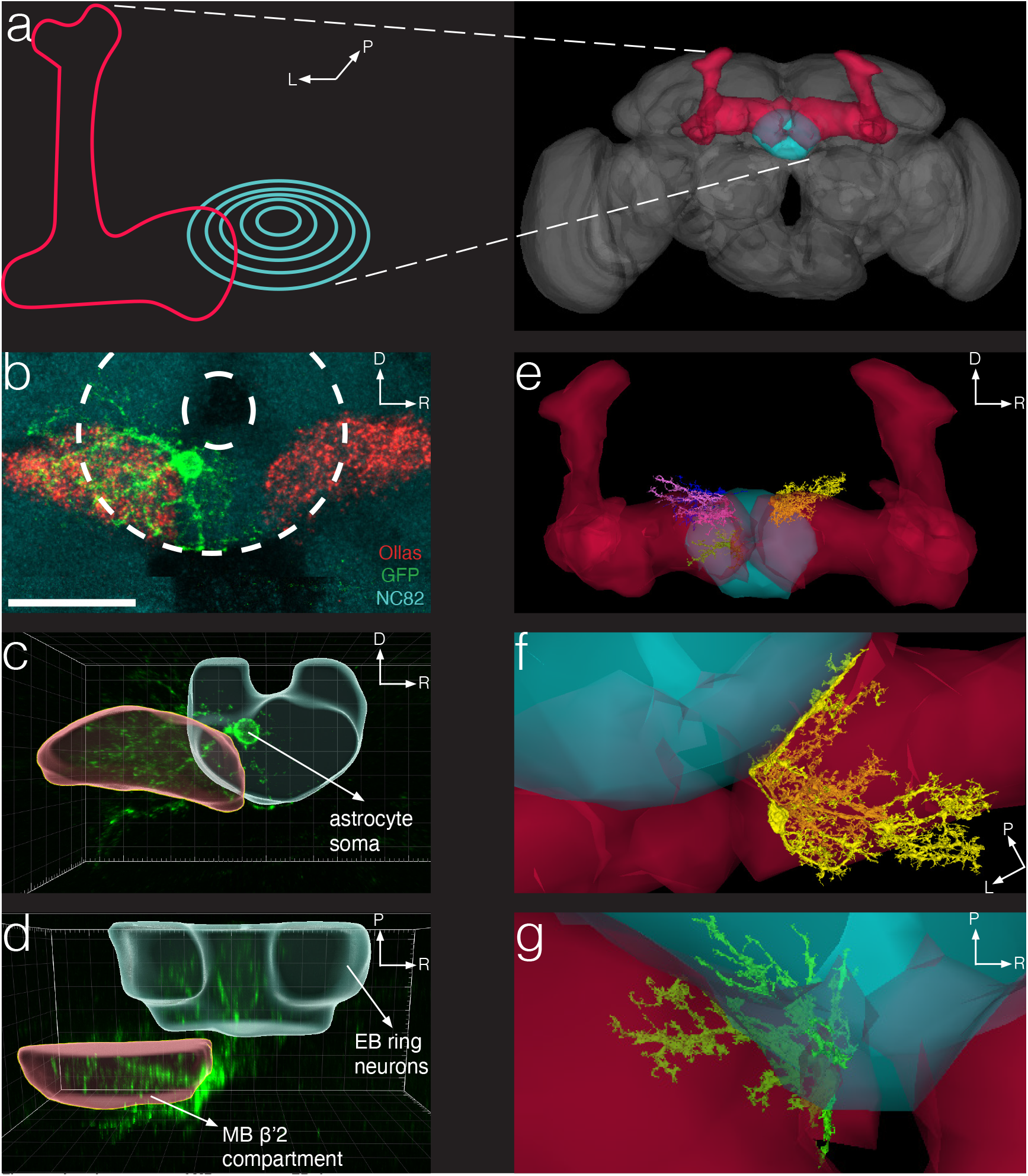
Local astrocytes of MB connect to EB ring neurons. **a**. Illustration of MB (red) and EB (turquoise) positions, zoomed in (left) and in relation to whole fly brain (right). Right image was extracted from flywire.ai. **b**. Max intensity projection of MB β’2 compartment and EB ring structures. Astrocyte labelled in green. Dashed line outlines EB ring neuron structure. Scale bar 20 μm. **c-d**. Two angles of 3D reconstruction showing how local astrocyte connects both structures. **e**. 4 cell IDs annotated as astrocytes from the flywire connectome connect MB and EB structures. **f-g**. Two example cell IDs showing astrocyte morphology.

Next, we asked whether there are any cells in the recently published connectome dataset [12,13] that were classified as astrocytes and connect the MB and EB. Currently, we identified four astrocytes that indeed connect these two regions. This number will likely increase as cell reconstructions are ongoing. In accordance with our immunostaining results, astrocyte cell bodies are located medially and attached to MB compartments with laterally projecting processes (**Fig. 4 e-g**).

## Discussion

Understanding the roles that astrocytes play in signal processing and animal behaviour requires targeting specific astrocyte subpopulations. In this study, we examined and optimized the genetic tool TRACT to label local astrocytes associated with user-defined *Drosophila* neuronal circuits in adults. The success of astro-TRACT relies on sufficient synaptic density. Higher synaptic density driven by the neuronal driver maximizes the success rate of labeling local astrocytes and minimizes unspecific neuronal labeling. The labelling sensitivity may vary between different neuronal circuits, as we have observed that astrocytes are labelled consistently for certain cell types (MB) regardless of the driver used, but not for others (EB-R5) (**Fig. S1**). To enhance labelling specificity, we switched the binary systems of TRACT, allowing its combination with highly specific neuronal split-Gal4 drivers. Furthermore, the extensive Gal4 driver collection creates high flexibility for interested users. In addition, in the future astro-TRACT can be adapted to other glial cell classes like ensheathing and perineurial glia by replacing the astrocyte enhancer, providing versatility in labeling experiments.

The tool’s greatest flexibility is provided by its plug-in design of astrocyte effectors. The available library of effectors under UAS control is vast, including calcium activity indicators, gene knockdown with RNAi and CRISPR-Cas9 for functional studies and GFP or other tag reporters for morphology characterization. The combination of these available tools with astro-TRACT allows users to adapt the tool for their specific research purposes. Furthermore, the labelling strategy is based on physical proximity of astrocyte to synapse rather than astrocyte-specific secretion molecules, making it a valuable tool for studying unknown astrocyte-neuron signaling mechanisms. By using functional indicators under UAS control in local astrocytes, astro-TRACT provides opportunities to unravel unknown astrocyte-neuron signaling pathways and to explore how these interactions may vary across different brain regions.

There are two alternatives to label individual fly astrocytes: single clone labeling with MARCM (Mosaic analysis with a repressible cell marker) [17] and MCFO (multi-color Flp out) [16]. In contrast to astro-TRACT, MARCM labels cells randomly and inconsistently between experiments. The multi-color mosaic technique MCFO can only label a cell type as specific as the driver used with it. In case of astrocytes, this means it will be expressed in astrocytes across the brain, which complicates analyses of individual astrocytes. Astro-TRACT provides higher reproducibility and specificity compared to these two existing tools, because local astrocytes of a chosen specific neuronal circuit are labelled.

Here, we used astro-TRACT to examine astrocyte projections from MB compartments to the close-by EB region. We found that astrocytes that surround compartments γ4, γ5, β’2, β2 connect to the EB. We also identified similarly positioned cells in the FlyWire connectome. We propose that this MB-EB connection by astrocytes may facilitate efficient communication between these regions. As the computational center for learning and memory in the fly brain, the MB processes sensory and cognitive information [18]. Its activity is accompanied by an accumulation of sleep need, indicated by increases of pre-synaptic proteins in response to sleep deprivation [19]. Therefore, astrocytes connecting MB to EB may transfer sleep pressure that accumulates in an experience and activity-dependent manner in the MB [20] to the sleep drive regulating R5 neurons of the EB [21].

In summary, we adapted the genetic tool TRACT to be used with specific split-Gal4 drivers targeting local astrocytes and characterized their expression specificity, sensitivity and reproducibility in the fly MB. In the future, this tool can provide the resolution necessary to identify novel signal transmission mechanisms between astrocytes and neurons.

## Methods

### Generation of *Drosophila* lines

#### *alrm* (or R86E01-DSCP)-nlg-synthetic_Notch-anti_CD19-Gal4-V5

The enhancer *nSyb* of the nSyb-nlg-SNTG4-V5 construct (gift from Lois lab, Caltech) was cut out with Nhe1 and EcoR1. Two lines, one with the 5069 bp long *alrm* enhancer [22] and another one with the 3140 + 155 bp R86E01-DSCP enhancer (Flylight) were constructed by cloning the respective enhancer fragment with overlapping homology arms into the nlg-SNTG4-V5 backbone by Gibson assembly (NEB). The following primers were used for PCR amplification of the enhancers:

*alrm*-F 5’ – GGA ACT AGG CTA GCA CTA CGC ACA GAT GTG GTC ATC TGA ATA GG – 3’;

*alrm*-R 5’ – CAC CCA TGG TGG AAT TCT AGT GGC GAT CCT TTC GCT CGG GAG C – 3’;

R86E01-F 5’ – GAA AAT GCT TGG ATT TCA CTG GAA CTA GGC TAG CCA CAT AAT ACT CTA CAG GGC ATC CAC – 3’;

R86E01-R 5’ – GAT CCC CGG GCG AGC TCG GAT TGT GAA GCC CCA AGA GTA C – 3’;

DSCP-F 5’ – GTA CTC TTG GGG CTT CAC AAT CCG AGC TCG CCC GGG GAT CG – 3’; DSCP-R 5’ – CCC AGG AGC TGG GTA GGG ACA CCC ATG GTG GAA TTC GTT TGG TAT GCG TCT TGT GAT TC – 3’

#### *alrm* (or R86E01-DSCP)-nlg-synthetic_Notch-anti_CD19-QF2

Another two lines were generated to contain the QF2 transcription factor instead of Gal4. The transcription factor QF2 was isolated from pCasper4-QF#7-hsp70 (Addgene #46135). The backbone *alrm* (or R86E01-DSCP)-nlg-synthetic_Notch-anti_CD19 was cut with Aat2 and Asc1. The 530 bp fragment of the part of the nlg-synthetic_Notch-anti_CD19 that was cut away from the backbone was amplified from nSyb-nlg-SNTG4-V5 and assembled with the amplified QF2 transcription factor with overlapping homology between inserts and backbone by Gibson assembly (NEB). The following primers were used for PCR amplification of QF2 and the 530 bp fragment of nlg-synthetic_Notch-anti_CD19:

QF2-F 5’ – GTA CAT ATT CAG GAA ATC TCA GTC GGC AGC TCT AGA CCA CCC AAG CGC AAA ACG CTT AAC – 3’;

QF2-R 5’ – CTT TAG TCG ACG GTA TCG ATA GAC GGG CGC GCC TCA TCA CTG TTC GTA TGT ATT AAT GTC – 3’;

nlg-synthetic_Notch-anti_CD19-F 5’ – GGT GGA ACT GCC ACC ATC ATC GTC GAC GGG CCA GAC GTC – 3’;

nlg-synthetic_Notch-anti_CD19-R 5’ – GCT TGG GTG GTC TAG AGC TGC CGA CTG AGA TTT CCT GAA TAT GTA CAC GTT TCT TGC CGC – 3’.

#### 10xUAS-IVS-nSyb-CD19

The 13xLexAop2 fragment of the construct 13xLexAop2-IVS-nsyb::CD19 (gift from Carlos Lois, Caltech) was removed and replaced by the 10xUAS fragment isolated from pJFRC81-10xUAS-IVS-Syn21-GFP-P10 (Addgene #36432) with Bgl2 and Hind3 by conventional T4 DNA ligase-mediated cloning (NEB).

The transgenes were injected into fly embryos in-house in the iso31 background by site-directed PhiC31-mediated insertion into VK00027 (*alrm*-nlg-synthetic_Notch-anti_CD19-Gal4-V5, *alrm*-nlg-synthetic_Notch-anti_CD19-QF2-V5, R86E01-DSCP-nlg-synthetic_Notch-anti_CD19-Gal4-V5) and JK73A (10xUAS-IVS-nSyb-CD19-Ollas) on the third chromosome and JK22C (*alrm*-nlg-synthetic_Notch-anti_CD19-QF2-V5, R86E01-DSCP-nlg-synthetic_Notch-anti_CD19-QF2-V5, 10xUAS-IVS-nSyb-CD19-Ollas) on the second chromosome.

After injection, one of multiple transformants was selected for the highest labelling specificity and efficiency. The chosen transformant was outcrossed 5 times to iso31. Subsequently, multiple genetic recombinations were performed. Various combinations of transgenes and injection sites were recombined to optimize the labelling efficiency and specificity.

#### Multi-color Flp-Out (MCFO) analyses

MCFO is a technique in which three different tags under UAS control are by default silenced by FRT-flanked transcriptional terminators [16]. Heat-shock induced FLPase expression randomly removes the terminators in individual cells that also express a GAL4 driver. This results in a mosaic of differently colored cells within the user-defined cell type. Flies with genotype HsFlpG5.Pest; alrm-synthetic_Notch-anti_CD19-Gal4; UAS-McFlip (UAS-STOP-smGFP-HA, UAS-STOP-smGFP-V5, UAS-STOP-smGFP-FLAG), MB247-LexA, LexAop-nSyb-CD19-Ollas were raised at 18°C. Adult 2-4 days-old flies were heat-shocked by being placed into a 37°C water bath for 5 minutes. Subsequently, flies recovered for 24 hours at 25°C before dissection.

#### Immunohistochemistry on whole-mount adult *Drosophila* brain GFP, Ollas tag and NC82

After dissection in S2 medium, fly brains were fixed in 2% paraformaldehyde for 55 minutes on a nutator. Next, brains were washed nutating at least four times for 15 minutes each in 0.5% PBST. Following that, the brains were blocked for 1.5 hours at room temperature in 5% Normal Goat Serum [Jackson Immuno Research 005-000-121] in 0.5% PBST. Then they were incubated in primary antibodies overnight at 4°C (Rabbit anti GFP 1:500 [Invitrogen A11122], Rat anti Ollas 1:200 [Novus Biologicals NBP1-06713], Mouse anti NC82 1:25 [DSHB]). The next day, brains were washed at least four times for 15 minutes each in 0.5% PBST. The secondary antibody (anti-Rabbit 488 1:1000 [Invitrogen A11008], anti-Rat 568 1:500 [Invitrogen A11077], anti-Mouse 647 1:300 [Invitrogen A21235]) incubation followed overnight at 4°C. Finally, the brains were washed at least four times for 15 minutes each in 0.5% PBST and left to incubate overnight at 4°C in Vectashield before mounting.

#### MCFO

The protocol for mosaic labelling was based on the ‘FlyLight Protocol – MCFO IHC for Adult Drosophila CNS’ and followed the same protocol as above with the following exceptions: The primary (Rat anti Flag 1:200 [Novus Biologicals NBP1-06712], Rabbit anti HA 1:300 [Roche 11867423001], Mouse anti NC82 1:25) and secondary antibodies (anti-Mouse 488 1:400 [Invitrogen A11029], anti-Rabbit 594 1:500 [Jackson Immuno Research 711-585-152], anti-Rat 647 1:300 [Invitrogen A21247]) were incubated for four hours at room temperature before 2 or 3-4 overnights at 4°C for primary and secondary, respectively. After the post-secondary antibody washes, an incubation in 5% Normal Mouse Serum (NMS) [Jackson Immuno Research 015-000-120] in 0.5% PBST followed for 1.5 hours on a nutator. Subsequently, brains were incubated in direct label DL550 Mouse anti-V5 antibody (1:500) [BioRad MCA1360D550GA] in 5% NMS in 0.5% PBST for 4 hours at room temperature and another one overnight at 4°C. Finally, brains were washed at least 4 times for 15 minutes each in 0.5% PBST at room temperature and left to incubate overnight at 4°C in Vectashield.

#### Image acquisition and analysis

Mounted brains were imaged using either a Zeiss Airyscan 880 or a Nikon NiE A1R confocal microscope. Images were processed in Fiji ImageJ (NIH). 3D reconstruction, segmentation and visualizations were generated with Imaris (Bitplane, version 10.0.1).

#### Connectome data analyses

Estimates of synaptic density by driver line were analyzed using the natverse libraries (v 1.10.4) [23] in R (v 4.3.3). Cell IDs in the hemibrain:v1.2.1 dataset were matched by driver with the neuronbridge_line_contents function of the neuronbridger package (v 2.2.0). For 11 of the 15 MB-projecting drivers, we were able to map connectome cell IDs. We then extracted the number of pre-synapses per MB compartment per cell ID and averaged this number across all cell IDs by driver. Then we multiplied this average to the average number of cells of the cell type targeted by the driver [18] arriving at an estimated number of pre-synapses per compartment per driver. Plotting the estimated synapse number by MB compartment was performed with ggplot2 (v 3.5.1).

To visualize putative astrocytes connecting MB and EB in the connectome, we scanned 141 cells flagged as ‘not_a_neuron’ and centrally located in the brain that are listed in the Supplemental Information of [12]. We identified 4 such cell_IDs (720575940617224498, 720575940607160753, 720575940627370277, 720575940648849540). 3D cell visualizations were directly extracted from codex.flywire.ai.

## Acknowledgements

We thank members of the Liu lab for discussions. Imaging was supported by the light microscopy expertise unit at the VIB-KU Leuven Center for Brain & Disease Research. We thank Carlos Lois (Caltech) for sharing plasmids and fly stocks. We also thank the Bloomington *Drosophila* Stock Center for providing fly stocks used in this study. We thank the Princeton FlyWire team and members of the Murthy and Seung labs, as well as members of the Allen Institute for Brain Science, for the development and maintenance of FlyWire (supported by BRAIN Initiative grants MH117815 and NS126935 to Murthy and Seung). We also acknowledge members of the Princeton FlyWire team and the FlyWire consortium for neuron proofreading and annotation. Special thanks to Marissa Sorek from the FlyWire team for support and Nikitas Serafetinidis, Joseph Hsu, Arti Yadav and Ryan Willie from the FlyWire Consortium for contributing >10% of the edits of cell reconstructions with cell_IDs 720575940617224498, 720575940607160753, 720575940627370277, 720575940648849540. This work was funded by a starting grant of the European Research Council to S.L. (#758580). J.D. held a PhD Fellowship of the Research Foundation - Flanders (FWO, 11D8820N and 11D8822N).

## Author Contributions

J.D. and S.L. conceived the study; J.D. performed molecular cloning; L.N. and J.Y. performed construct injections into fly embryos; J.D., S.M. and F.H. performed immunohistochemistry and imaging; J.D. performed image processing and analysis; J.D. performed computational analyses; J.D. and S.L. wrote and revised the manuscript.

