## Supplementary material for "Genetic targeting of astrocytes associated with specific neuronal circuit in adult *Drosophila*": Figure S1

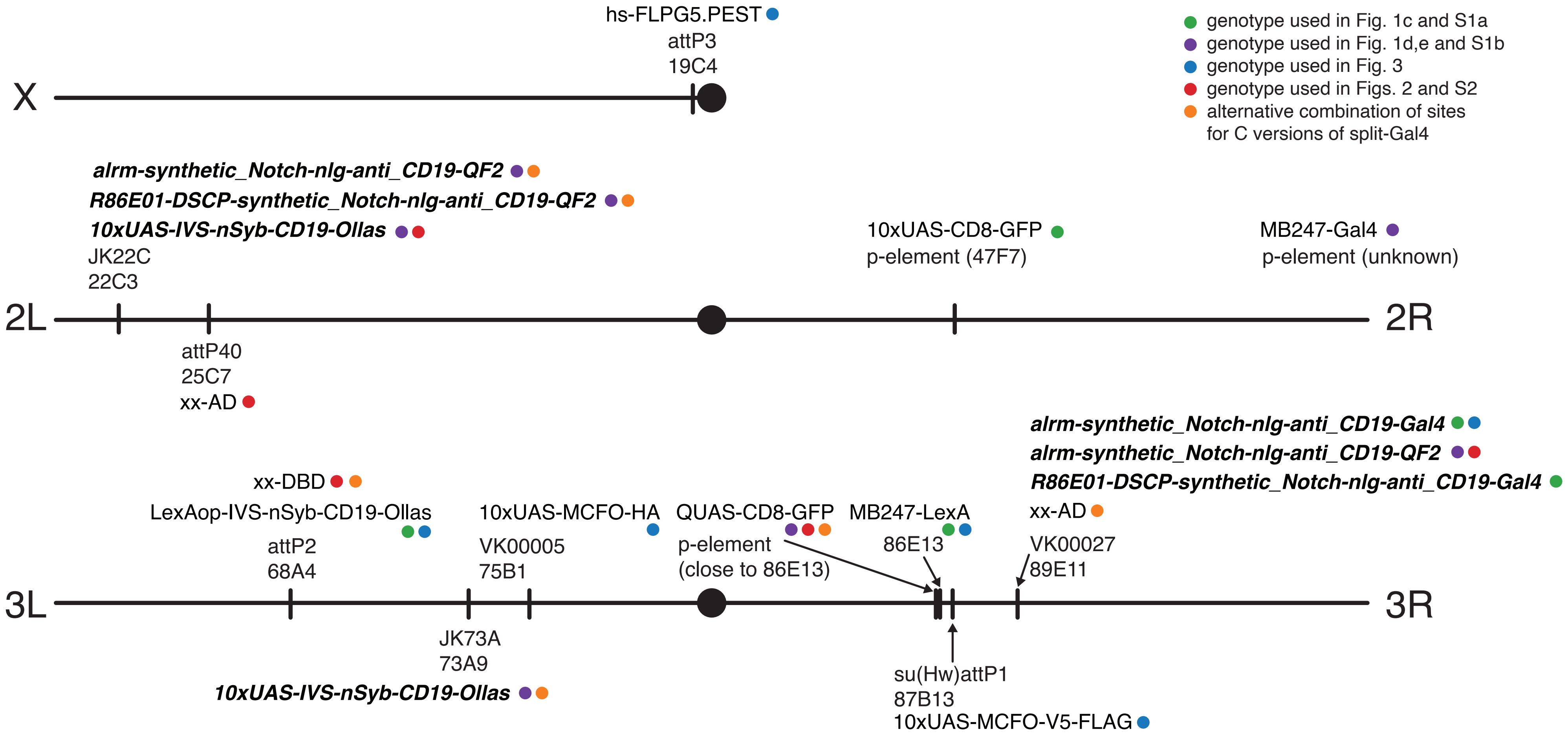


**Figure S1. Genetic elements used in present study shown by injection site in three chromosomes of *Drosophila* genome.**
Red elements belong to genotype used in Figure 2 and S2. All split-Gal4 (combination of AD and DBD) are inserted in attP40 (AD) and attP2 (DBD) with the exception of MB315C, MB112C, MB320C (AD in VK00027, DBD in attP2). Combination of genetic elements that can be used for those lines to avoid conflict of elements at the same site are shown in pink.
Blue elements belong to genotype used in Figure 3.
Bolded elements were generated in this study.
