## Supplementary material for "Genetic targeting of astrocytes associated with specific neuronal circuit in adult *Drosophila*": Figure S2

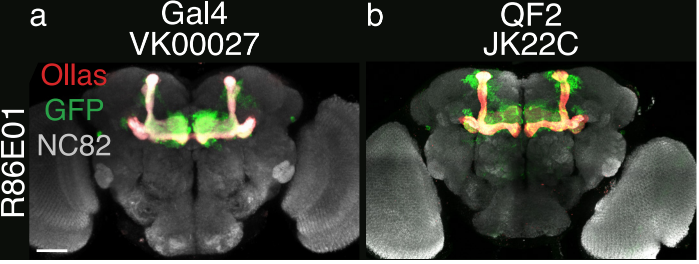


**Figure S2. Expression of astrocytes around the Mushroom Body, with R86E01 enhancer.**

**a.** Gal4 version injected in VK00027.

**b.** QF2 version injected in JK22C.
