## Supplementary material for "Genetic targeting of astrocytes associated with specific neuronal circuit in adult *Drosophila*": Figure S3

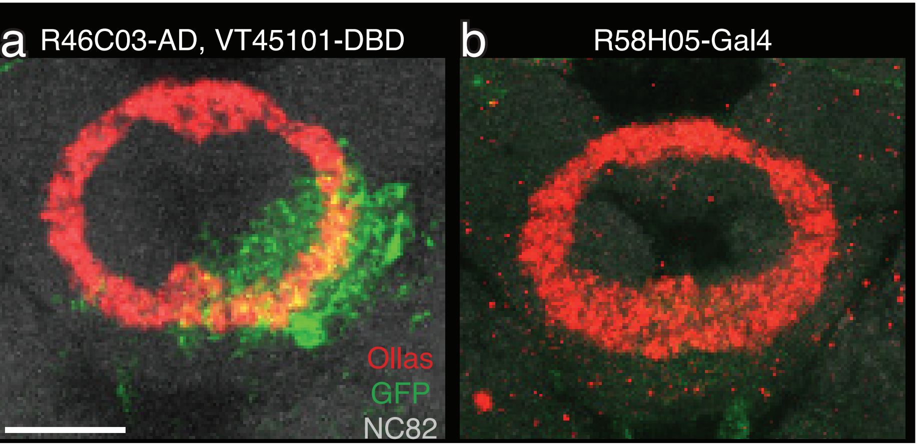


**Figure S3. Local astrocytes of EB R5 neurons.**
**a.** Sparse labelling with split-Gal4 driver R46C03-AD, VT45101-DBD. Scale bar 50 µm.
**b.** No astrocyte labelling with R58H05-Gal4.
