## Supplemental Tables for "Genetic targeting of astrocytes associated with specific neuronal circuit in adult *Drosophila*"

**Table S1. Labelling results of astro-TRACT by MB driver line**

| split-Gal4 driver | labels | cell # | individual astro | brains labelled (%) | unspecific neurons |
|---|---|---|---|---|---|
| MB441B | PAM-γ3 | 9-23 | yes | 100 | yes |
| MB042B | PAM-y3,4,5 | | yes | 100 | yes |
| MB196B | PAM-y3,4,5 | | yes | 100 | minor |
| MB315C | PAM-y5 | 8-21 | yes | 100 | yes |
| MB312B | PAM-γ4 | 13-17 | yes | 50 | yes |
| MB607B | KC γd | 75 | yes | 80 | yes |
| MB009B | KC y | | yes | 66,66666667 | yes |
| MB112C | MBON-y1 | 1 | no | NA | yes |
| MB298B | MBON-γ4>γ1γ2 | 1 | no | NA | yes |
| MB504B | PPL1-y1,2, α'1,2 | | no | NA | yes |
| MB320C | PPL1-y1 | | no | NA | yes |
| MB025B | PAM-β'1 | 13-14 | yes | 100 | no |
| MB188B | PAM-β'1 | | yes | 100 | yes |
| MB463B | KC α'/β' | 210 | yes | 100 | no |
| MB418B | KC α'/β'm | 140 | yes | 100 | yes |
| MB185B | KC α/βs | 500 | yes | 100 | no |
| MB008B | KC α/β | | yes | 100 | minor |
| MB371B | KC α/βp | 90 | yes | 50 | yes |

| connects to EB (%) | astro labelling notes |
|---|---|
| 0 | y3 and weak pre-synaptic signal |
| 100 | y4,5 |
| 25 | y4,5 |
| unclear | |
| unclear | |
| 0 | |
| no, anterior to EB | only one astro each |
| NA | |
| NA | |
| NA | |
| NA | |
| 0 | |
| 0 | astro labelling with very weak pre-synaptic signal |
| 100 | |
| 100 | |
| 100 | |
| 66,66666667 | along the lobe |
| 0 | typically not medial |

**Table S2. Synapse numbers by MB driver line based on public connectomics data**

| ROI | line | individual_astı | ct | ncell | roipre_avg | n_matched_ne |
|---|---|---|---|---|---|---|
| a'1 | MB418B | yes | KC | 140 | 33 | 1 |
| a'2 | MB418B | yes | KC | 140 | 28 | 1 |
| a'3 | MB418B | yes | KC | 140 | 29 | 1 |
| a1 | MB112C | no | MBON | 1 | 34,5 | 2 |
| a1 | MB504B | no | PPL1 | 12 | 53 | 1 |
| a2 | MB112C | no | MBON | 1 | 37,5 | 2 |
| a2 | MB504B | no | PPL1 | 12 | 64 | 1 |
| a3 | MB112C | no | MBON | 1 | 31,5 | 2 |
| a3 | MB504B | no | PPL1 | 12 | 60 | 1 |
| b'1 | MB025B | yes | PAM | 13 | 112,1578947 | 19 |
| b'1 | MB418B | yes | KC | 140 | 43 | 1 |
| b'2 | MB418B | yes | KC | 140 | 87 | 1 |
| b1 | MB112C | no | MBON | 1 | 25,5 | 2 |
| b1 | MB504B | no | PPL1 | 12 | 29 | 1 |
| b2 | MB112C | no | MBON | 1 | 24 | 2 |
| b2 | MB504B | no | PPL1 | 12 | 36 | 1 |
| g1 | MB298B | no | MBON | 1 | 170 | 2 |
| g1 | MB607B | yes | KC | 75 | 88 | 1 |
| g2 | MB298B | no | MBON | 1 | 115 | 2 |
| g2 | MB607B | yes | KC | 75 | 51 | 1 |
| g3 | MB188B | yes | PAM | 13 | 59,4 | 5 |
| g3 | MB441B | yes | PAM | 16 | 58,14285714 | 14 |
| g3 | MB607B | yes | KC | 75 | 69 | 1 |
| g4 | MB196B | yes | PAM | 15 | 124 | 1 |
| g4 | MB312B | yes | PAM | 15 | 82 | 23 |
| g4 | MB607B | yes | KC | 75 | 212 | 1 |
| g5 | MB315C | yes | PAM | 14 | 67,04651163 | 43 |
| g5 | MB607B | yes | KC | 75 | 51 | 1 |

| avg_n |
| ---: |
| 4620 |
| 3920 |
| 4060 |
| 34,5 |
| 636 |
| 37,5 |
| 768 |
| 31,5 |
| 720 |
| 1458,052632 |
| 6020 |
| 12180 |
| 25,5 |
| 348 |
| 24 |
| 432 |
| 170 |
| 6600 |
| 115 |
| 3825 |
| 772,2 |
| 930,2857143 |
| 5175 |
| 1860 |
| 1230 |
| 15900 |
| 938,6511628 |
| 3825 |
